# A microRNA that controls the emergence of embryonic movement

**DOI:** 10.1101/2024.01.24.577117

**Authors:** Jonathan A. C. Menzies, Andre M. Chagas, Tom Baden, Claudio R. Alonso

**Affiliations:** Department of Neuroscience, Sussex Neuroscience, School of Life Sciences, University of Sussex, Brighton BN1 9QG, UK

**Keywords:** Drosophila, embryo, movement, microRNA, neuron

## Abstract

Movement is a key feature of animal systems, yet its embryonic origins are not fully understood. Here we investigate the genetic basis underlying the embryonic onset of movement in *Drosophila* focusing on the role played by small non-coding RNAs (microRNAs, miRNAs). To this end, we first develop a quantitative behavioural pipeline capable of tracking embryonic movement in large populations of fly embryos, and using this system, discover that the *Drosophila* miRNA *miR-2b-1* plays a role in the emergence of movement. Through the combination of spectral analysis of embryonic motor patterns, cell sorting and RNA *in situs*, genetic reconstitution tests, and neural optical imaging we define that *miR-2b-1* influences the emergence of embryonic movement by exerting actions in the developing nervous system. Furthermore, through the combination of bioinformatics coupled to genetic manipulation of miRNA expression and phenocopy tests we identify a previously uncharacterised (but evolutionarily conserved) chloride channel encoding gene – which we term *<u>Mo</u>vement Modula<u>tor</u>* (*Motor)* – as a genetic target that mechanistically links *miR-2b-1* to the onset of movement. Cell-specific genetic reconstitution of *miR-2b-1* expression in a null miRNA mutant background, followed by behavioural assays and target gene analyses, suggest that *miR-2b-1* affects the emergence of movement through effects in sensory elements of the embryonic circuitry, rather than in the motor domain. Our work thus reports the first miRNA system capable of regulating embryonic movement, suggesting that other miRNAs are likely to play a role in this key developmental process in *Drosophila* as well as in other species.

## RESULTS AND DISCUSSION

### A miRNA that impacts larval movement

Movement is the main output of the nervous system allowing animals to walk, fly, crawl, swim and maintain their posture, so that they can find prey, mate partners, escape predators and relocate within habitats. Despite the key biological and adaptive roles of movement across animal systems, how developing embryos manage to organise the necessary molecular, cellular, and physiological processes to initiate patterned movement is still unknown. Although it is clear that the genetic system plays a role, how genes control the formation, maturation and function of the cellular networks underlying the emergence of motor control remains poorly understood.

Recent work in our laboratory has shown that miRNAs – which are short regulatory non-coding RNAs that repress the expression of target genes [1, 2] – have pervasive roles in the articulation of complex movement sequences such as those involved in body posture control in the young *Drosophila* larva [3–6]; these observations, as well as those from others in *Drosophila* and other systems [7–11] hinted at the possibility that these non-coding RNA molecules might also be involved in the control of more fundamental aspects of motor development and control. To explore this question, we first searched for miRNAs that might affect the simple locomotor patterns of the *Drosophila* first instar larva. The L1 larva which is a convenient model to investigate the genetics of movement given that: (i) the assembly of the machinery for movement must be fully completed by this developmental stage so as to satisfactorily propel the animal into the external world, and (ii) if analysed sufficiently early, for example during the first few minutes after hatching, the animal had no real chance of compensating or learning ways around a putative defect, increasing the probability of detection by means of a suitable motor test. To extract a signature of larval movement, we applied a whole animal imaging method based on frustrated total internal reflection (FTIR) [12, 13] which renders high resolution and high contrast movies to both normal and miRNA mutant first instar larvae. This led us to discover that a single miRNA, *miR-2b-1*, had a significant impact on larval movement (Figure 1B-1C). *miR-2b-1* belongs to the *miR-2* family [14] and *ΔmiR-2b-1* mutant larvae show a substantial decrease in larval speed (Figure 1C) suggesting that absence of this specific genetic component compromises the ability of the larva to move normally. Given that these larval tests were conducted within a 30-min period post-hatching, we considered the possibility that the defects observed in the larvae stemmed from changes in earlier ontogenetic processes that occur in the embryo. More specifically, we decided to test the possibility that early embryonic movement patterns might be affected by the lack of normal expression of *miR-2b-1*.

**Figure 1.**
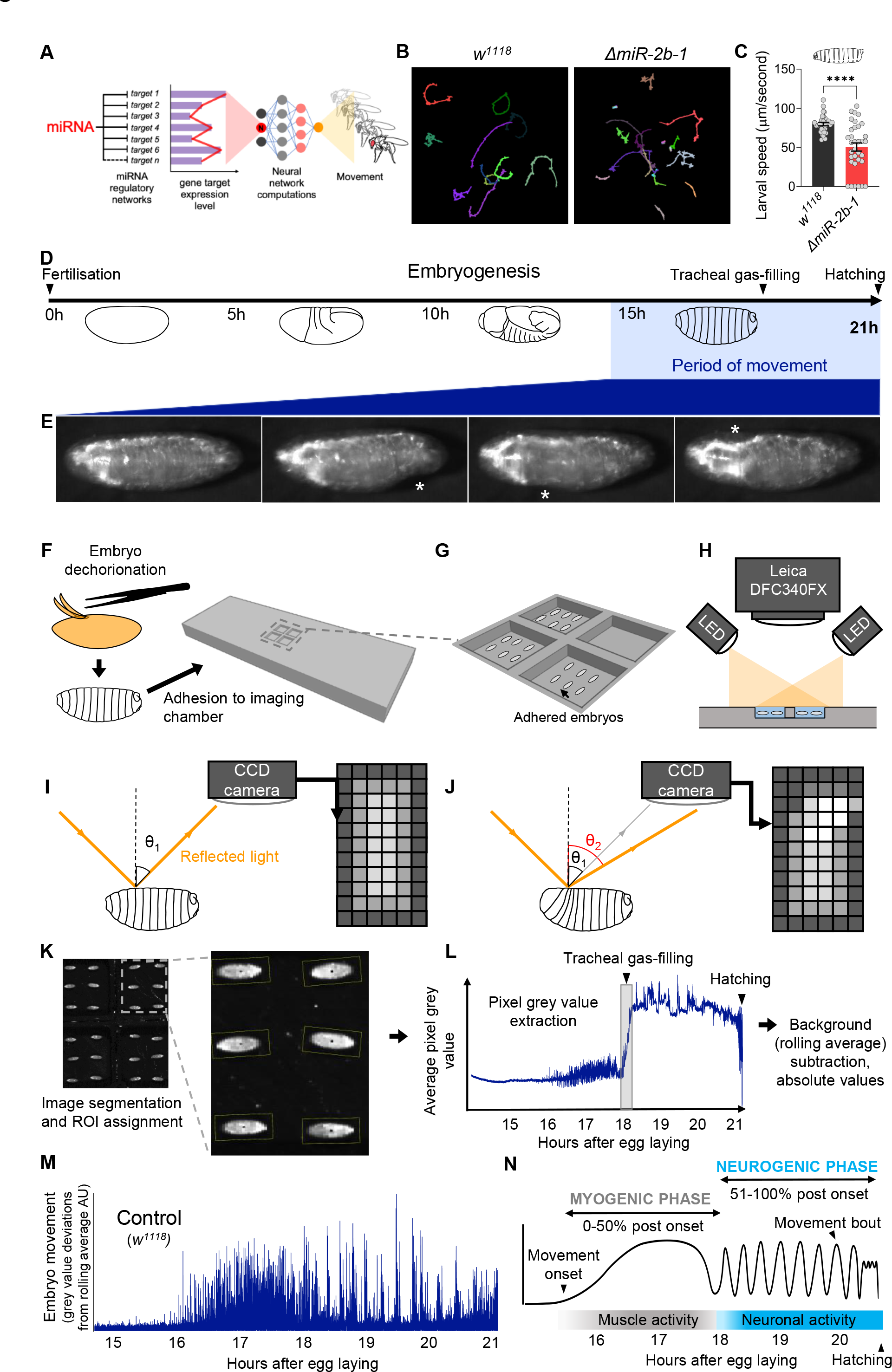
A novel approach for the quantification of embryonic movement (A) Diagram illustrating the point of action of miRNAs in the neural networks controlling behaviour. (B) Larval movement tracks for *w^1118^* (left) and *ΔmiR-2b-1* (right) larvae obtained using the frustrated total internal reflection based imaging system (FIM). (**C**) Quantification of average larval speed for *w^1118^* and *ΔmiR-2b-1* larvae using the FIM system. (**D**) Schematic describing the timeline of *Drosophila* embryonic development with the period of movement highlighted in blue. (**E**) Microscope images of a *Drosophila* embryo performing characteristic early movements, highlighted with asterisks. (**F-H**) Experimental pipeline for recording embryo movements. (F) eggshell removal (dechorionation) and adhesion to the imaging chamber, (G) imaging chamber design and (H) imaging set up under incident LED illumination. (**I-J**) Schematic describing the basis for movement detection: light is reflected from the embryo surface and internal structures and detected by a CCD sensor to generate a pixel map of the embryo (I). Embryonic movement changes the angle of reflected light, resulting in a different pixel map (J). (**K-N**) Pipeline for the quantification of embryonic movement. (K) representative image of the embryo movement chamber showing the assignment of regions of interest (ROIs) to individual embryos; (L) extraction of the mean grey value (MGV) for each frame (done in parallel for each ROI) allows the generation of raw movement traces for each individual embryo. Key developmental events that impact MGV (tracheal gas-filling, hatching) are indicated by arrowheads. (M) subtraction of the trace background calculated by rolling average removes slow changes in MGV that result from developmental events, allowing accurate quantification of deviations in MGV from baseline which represent movement over time (absolute values); (N) schematic of an idealised wild-type (*w^1118^*) movement trace with putative phases indicated.

Previous work had provided an excellent first characterisation of the onset of embryonic movement patterns in wild type embryos [15, 16]; these early studies were based on the manual annotation of representative videotaped muscle contractions [16] or GFP-labelled muscle Z-lines analysed under spinning disc confocal microscopy [15]. Despite their attributes, these approaches were highly labour intensive and lacked the necessary throughput required to simultaneously analyse large numbers of embryos enabling a sensitive genetic screen. In consequence, we developed a new automated approach capable of quantifying movement in large populations of embryos.

### A high throughput approach to quantify movement in Drosophila embryos

To monitor the onset of embryonic movement, which, in normal embryos, occurs during the final third of embryogenesis (i.e. 14-16h after egg laying (AEL) [15, 16] [Figure 1D-1E] we developed a 3D printed chamber system capable of hosting multiple embryos submerged under a thin layer of halocarbon oil to ensure adequate oxygenation and hydration (Figure 1F-1G) compatible with digital imaging by a charge coupled device (CCD) camera (Figure 1H). The camera captures the reflected light after its physical contact with the embryo; in this setting, even a subtle movement performed by the embryo results in a change in the path of reflected light, leading to variations in signal intensity detected by individual pixels in the CCD sensor, allowing for an accurate measurement of embryonic movement (Figure 1I-1J). To extract quantitative movement information from individual embryos we applied an image segmentation protocol to define regions of interest (ROIs) corresponding to each embryo and collected pixel intensity values from all ROIs at 4 frames/sec (Figure 1K-1L). The data allow us to plot variations in average grey pixel intensity over embryonic time, which provide a quantitative signature of the ontogeny of movement in the individual embryo (Figure 1M). From this, we were able to observe distinct phases of movement that are consistent with previous data: namely, the onset of a phase of disorganised movements ∼16 hours after egg laying (hAEL) and its transition into a phase characterised by rhythmic bursts of activity and inactivity ∼18 hAEL [15, 16]. These phases have been termed as ‘myogenic’ and ‘neurogenic’ based on their respective dependence on neural input (Figure 1N) [15, 17–19] , and have been observed in *Drosophila*, as well as in other systems, including vertebrates [20–24] strongly suggesting that this is a general feature of motor development. See Movie S1 for a high-resolution recording of *Drosophila* embryo movement.

To establish whether the readings of motor activity at the neurogenic phase detected by our approach were indeed dependent on neural activity, we expressed the inwardly rectifying potassium channel (*Kir*) [25] in all embryonic neurons using the pan-neuronal driver *elav-Gal4* seeking to suppress action potentials across all embryonic neuronal types. The results of this experiment show that whilst movement patterns during the early chaotic phase remain unchanged by this treatment (Figure 2C, 2E), motor activity at the rhythmic phase is almost completely eliminated (Figure 2C, 2F) strongly indicating that the emergence of this latter phase depends on normal neural activity. This is in agreement with previous observations of the effects of embryonic neural activity inhibition by other methods [15, 18]. In addition, spectral analysis demonstrates that the movement frequencies that characterise the rhythmic phase (Figure 2B, 2G) do not emerge in *Kir* embryos (Figure 2D, 2G). Altogether these observations suggest that *miR-2b-1* might exert its roles on embryonic movement, at least in part, due to action within the developing embryonic nervous system.

**Figure 2.**
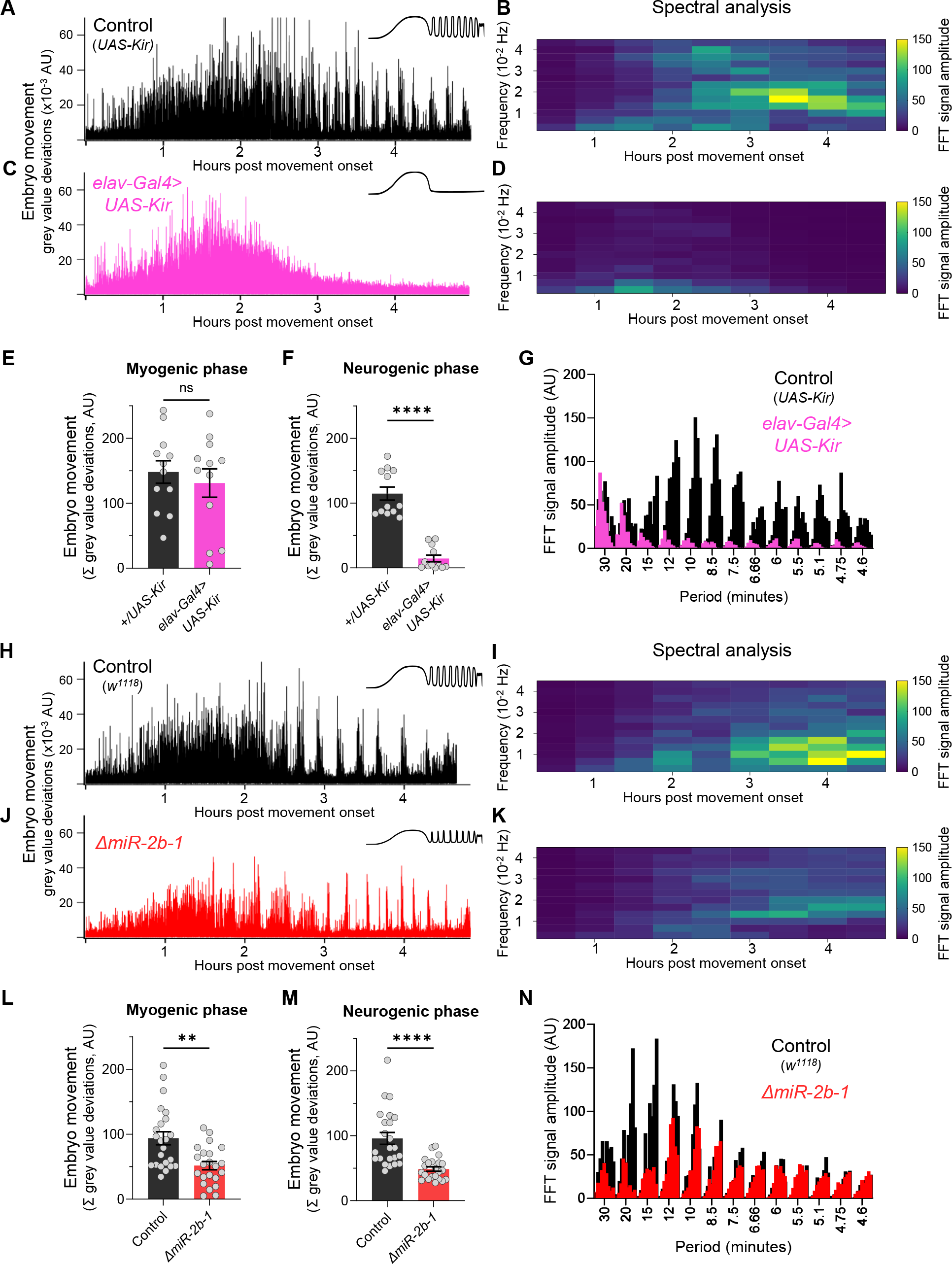
miR-2b-1 controls movements during the neurogenic phase of embryonic movement (**A**) Representative movement trace for control (*UAS-Kir*) animals. A concept diagram that summarises the pattern is shown in the top right. (**B**) Heat map showing the average frequency spectrogram for movements of control (*UAS-*Kir) animals, determined by fast fourier transform (FFT) analysis (1-hour sliding window with a discrete 30-min step size) from onset of embryonic movement to hatching. Brighter colours indicate a stronger amplitude of movement at a given frequency. (**C**) Representative movement trace for experimental (*Elav-Gal4>UAS-Kir*) animals. (**D**) Average frequency spectrogram for *Elav-Gal4>UAS-Kir* animals. (**E**) Summation of MGV deviations during the myogenic phase in control (*UAS-Kir*, black) and experimental (*Elav-Gal4>UAS-Kir*, pink) animals. (**F**) Summation of MGV deviations during the neurogenic phase in control (*UAS-Kir*, black) and experimental (*Elav-Gal4>UAS-Kir*, pink) animals. (**G**) Distribution of signal amplitudes across different movement periods (p) derived from the FFT frequency analysis shown in panels B and D. A higher signal amplitude is produced when more movement occurs with a particular periodicity. Bars of different height at each period sampled show data from individual embryos. (**H**) Representative movement trace for control (*w^1118^*) animals. (**I**) Heat map showing the average frequency spectrogram for movements of *w^1118^* control embryos. (**J**) Representative movement trace for experimental (*ΔmiR-2b-1*) animals. (**K**) Heat map showing the average frequency spectrogram for movements of *ΔmiR-2b-1* embryos. (**L**) Summation of MGV deviations during the myogenic phase in control (*w^1118^*, black) and experimental (*ΔmiR-2b-1*, red) animals. (**M**) Summation of MGV deviations during the neurogenic phase in control (*w^1118^*, black) and experimental (*ΔmiR-2b-1*, red) embryos. (**N**) Distribution of signal amplitudes across different movement periods derived from the FFT frequency analysis shown in panels I and K.

### The miRNA miR-2b-1 affects embryonic movement patterns

Given that the machinery for larval movement is assembled during embryogenesis [26–29] we considered the possibility that *miR-2b-1* might have an impact on the emergence of movement in the fly embryo. To explore this, we applied the approach described above to normal and *ΔmiR-2b-1* mutant embryos (Figure 2H, 2J). These experiments showed that *ΔmiR-2b-1* mutant embryos displayed a different pattern of embryonic movement when compared with their wild type counterparts. Although the distinct phases of embryonic movement are clearly recognisable in mutant embryos, the overall amount of movement appeared greatly reduced (Figure 2J).

Indeed, comparison of quantity of movement in wild type and *ΔmiR-2b-1* mutant embryos either during the earlier myogenic phase (Figure 2L) or during the neurogenic phase (Figure 2M) shows significantly lower levels of movement in mutants. Furthermore, spectral analyses of embryonic movement traces reveal that miRNA mutant embryos shift to a higher frequency of movement bouts during the rhythmic phase when compared to normal embryos (Figure 2I, 2K) and that average bout length is also shortened (Figure S1).

### miR-2b-1 expression and roles in the embryonic nervous system

The fact that removal of *miR-2b-1* impacts the neurogenic phase of embryonic movement suggests that this miRNA might be expressed in the nervous system and exert a functional role there. To further explore this model, we first conducted a spatial expression analysis in the developing embryo using fluorescence RNA *in situ* hybridisation (FISH). Taking advantage from the fact that *miR-2b-1* is located within the 3’ untranslated region (3’UTR) of the *Bruton tyrosine kinase* (*Btk*) gene [30, 31] (Figure 3A) we prepared an *in situ* probe to detect *Btk* transcripts. These FISH experiments show that the *miR-2b-1* precursor miRNA transcript (pre-miRNA) [2] is expressed in multiple embryonic tissues, including the CNS (Figure 3B). To further confirm the expression of *miR-2b-1* in the nervous system, we conducted fluorescence-activated cell sorting (FACS, Figure 3C) to isolate embryonic neuronal samples (labelled by means of *elav>GFP*) followed by RT-PCR for the mature miRNA transcript and observed *miR-2b-1* specific signal (Figure 3D). Therefore, the results of these two distinct and complementary methods provide strong evidence of neural expression of the miRNA. To gain more insight on the neural roles of *miR-2b-1* in regard to embryonic movement, we conducted a genetic reconstitution experiment in which we analysed the consequences of restoring *miR-2b-1* expression in the nervous system in an otherwise *ΔmiR-2b-1* null mutant (Figure 3E). Results in Figure 3F, S2A show that *elav*-driven expression of *miR-2b-1* in a *ΔmiR-2b-1* mutant background leads to a phenotypic rescue, producing embryos that display statistically indistinguishable movement patterns to those recorded in control embryos, indicating that neural expression of *miR-2b-1* is sufficient to restore a normal onset of embryonic movement. To further examine the biological roles of neural *miR-2b-1* expression, we assessed the impact of restoring miRNA expression in embryonic neurons on first instar larval locomotor patterns using the FIM approach described above (Figure 1B- C) and observed that when *ΔmiR-2b-1* mutant larvae are developmentally provided with pan-neuronal *miR-2b-1* expression, the characteristic miRNA larval mutant phenotype is rescued, with specimens moving at natural speed (Figure 3G, S2B). The experiments described above strongly indicate that expression of *miR-2b-1* in the nervous system is biologically relevant and sufficient to rescue the embryonic and larval movement defects observed in *ΔmiR-2b-1* mutants. They also suggest that the effects of *miR-2b-1* observed at earlier stages (myogenic phase) are possibly offset by normal neural expression of *miR-2b-1*.

**Figure 3.**
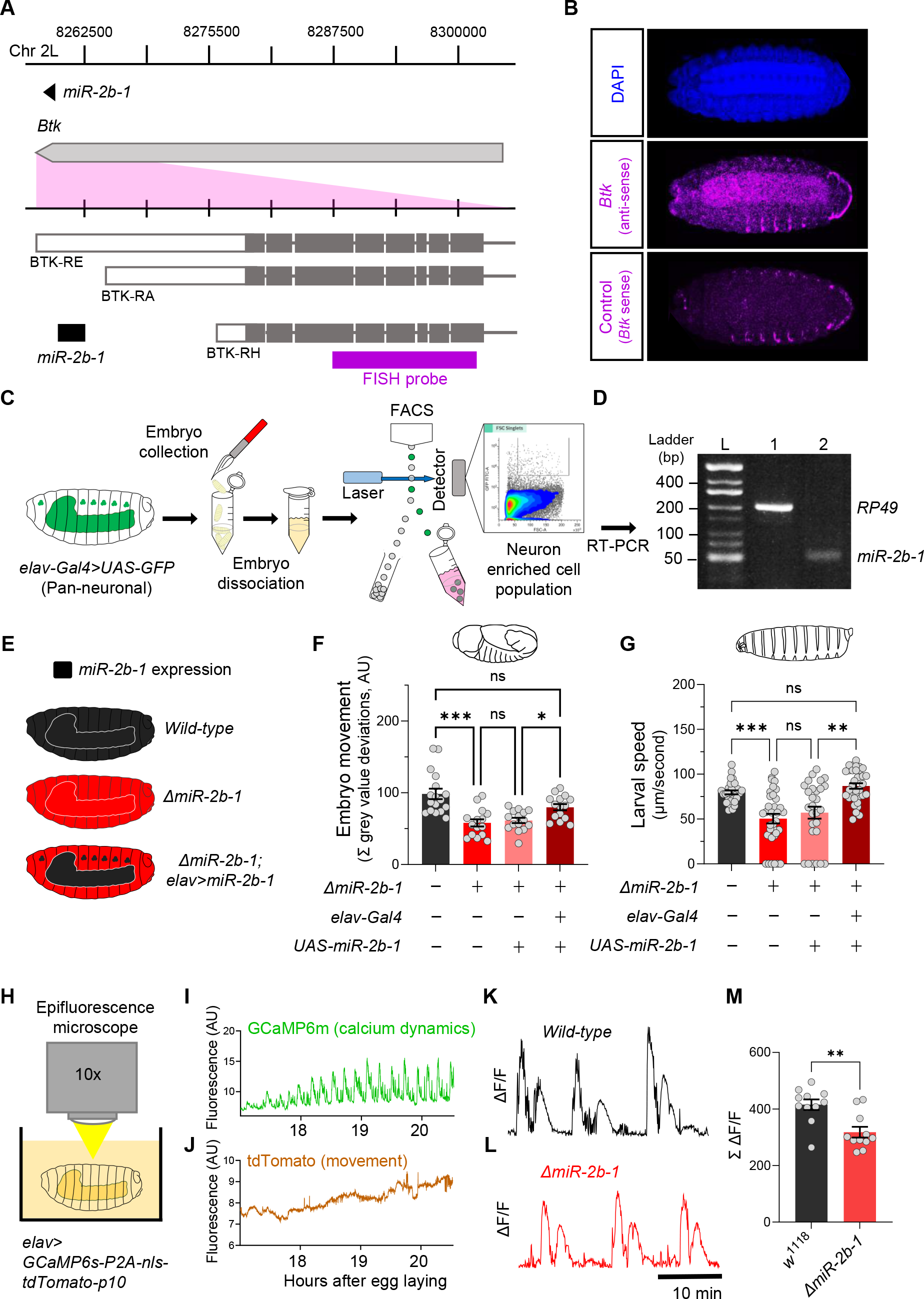
miR-2b-1 acts within neurons to regulate embryonic and larval movement (**A**) Gene diagram describing the *miR-2b-1* locus including the host gene *Btk* and its RNA transcripts. A fluorescence *in situ* hybridisation (FISH) probe for *Btk* is indicated by the magenta bar. (**B**) FISH experiment on control *w^1118^* embryos showing expression of the miRNA host *Btk* transcripts (antisense probe, middle panel) [DAPI stain in blue (upper image); *Btk* sense probe in magenta (lower image)]. (**C**) Experimental workflow for a fluorescence activated cell sorting (FACS) experiment. Neurons were labelled by *elav-Gal4>UAS-GFP* (left), followed by enzymatic and mechanical separation and isolation of GFP+ neurons. (**D**) RT-PCR analysis showing expression of mature *miR-2b-1* (right) detected in neurons at the onset of the neurogenic phase (*RP49* signal (left) shown as control). (**E**) Movement patterns were assessed in embryos with normal *miR-2b-1* expression (wild-type, black), null *miR-2b-1* mutants (red) and in mutant embryos in which *miR-2b-1* expression was restored (reconstituted) specifically in neurons (red and black). (**F**) Summation of MGV deviations during the neurogenic phase of embryonic movement in control *w^1118^* embryos (black bar); *ΔmiR-2b-1* mutant embryos (bright red bar); *ΔmiR-2b-*, *UAS-miR-2b-1* parental control embryos (faded red bar); *ΔmiR- 2b-1*, *Elav-Gal4>UAS-miR-2b-1* embryos (black and red lined bar). (**G**) Average L1 larval speed for the same genotypes used in the embryonic genetic reconstitution experiment. (**H**) Schematic describing the experimental setup for fluorescence imaging of *elav>GCaMP6s-P2A-nls-tdTomato- p10* embryos under an epifluorescence microscope. This design allows simultaneous detection of calcium dynamics (GCaMP6s) and movement (tdTomato). (**I**) Representative GCaMP6s trace (green) from control *w^1118^* embryos over the neurogenic phase of embryonic movement. (**J**) Representative tdTomato trace from the same embryo as in panel I, acting as a passive fluorescence reporter used to subtract changes in GCaMP6s signal induced by embryonic movement. (**K**) Representative ΔF/F trace for *w^1118^*embryos. (**L**) Representative ΔF/F trace for *ΔmiR-2b-1* mutant embryos. (**M**) Summation of ΔF/F signal during embryogenesis in control *w^1118^* embryos (black bar) and *ΔmiR-2b-1* mutant embryos (red bar).

In turn, this suggests that absence of *miR-2b-1* must impinge a morphological and/or a functional deficit in the developing nervous system of the embryo. To tease apart these potential biological effects, we examined the structure of the nervous system in normal and *ΔmiR-2b-1* mutant embryos by means of immunohistochemistry and confocal microscopy and observed no detectable differences (Figure S2C). In contrast, analysis of neural activity patterns in the embryo by means of GCaMP6 functional imaging using a movement distortion correction approach (i.e. tdTomato, Figure 3H-3J [17]), shows that miRNA mutant embryos have a reduced level of calcium dynamics when compared with their control counter parts (Figure 3K-3M); notably, this occurs during a previously identified ‘critical period’ when neural activity levels are of crucial importance to the development of stable neural circuits [32–34]. Consistently with this previous work, artificial reduction of embryonic neural activity via optogenetic control leads to a significant decrease in larval speed (Figure S4A-G).

### A genetic link between miR-2b-1 and embryonic movement

Our gene expression, genetic reconstitution, morphological and functional imaging data support a model in which *miR-2b-1* plays a physiological role in the developing embryonic nervous system. This raises the question of how might this regulatory miRNA system interact with the physiological control of the neuron during embryogenesis. To explore the genetic elements that link *miR-2b-1* to its role in embryonic movement we searched for candidate *miR-2b-1* target genes using the ComiR bioinformatic platform (Figure 4B) [35, 36]. A common issue with bioinformatic predictions of miRNA targets is the generation of false positives; in this regard, ComiR integrates multiple miRNA target prediction algorithms – each one with its intrinsic strengths and weaknesses[37–40] – seeking to identify a set of consistent *bona fide* miRNA targets that satisfy the filters of multiple algorithms, thus reducing the generation of false positives[35, 36]. Applying ComiR to *miR-2b-1* produced a list of high probability targets organised in the form of an ascending ranking (Figure 4B). At the very top of the list was *CG3638*, an uncharacterised *Drosophila* gene predicted to encode a chloride channel protein; this highly ranked target was of interest to us because of its potential role in the physiological control of anionic conductances, and its broad evolutionary conservation across insects and mammals[41], including humans (Figure 4E-I).

**Figure 4.**
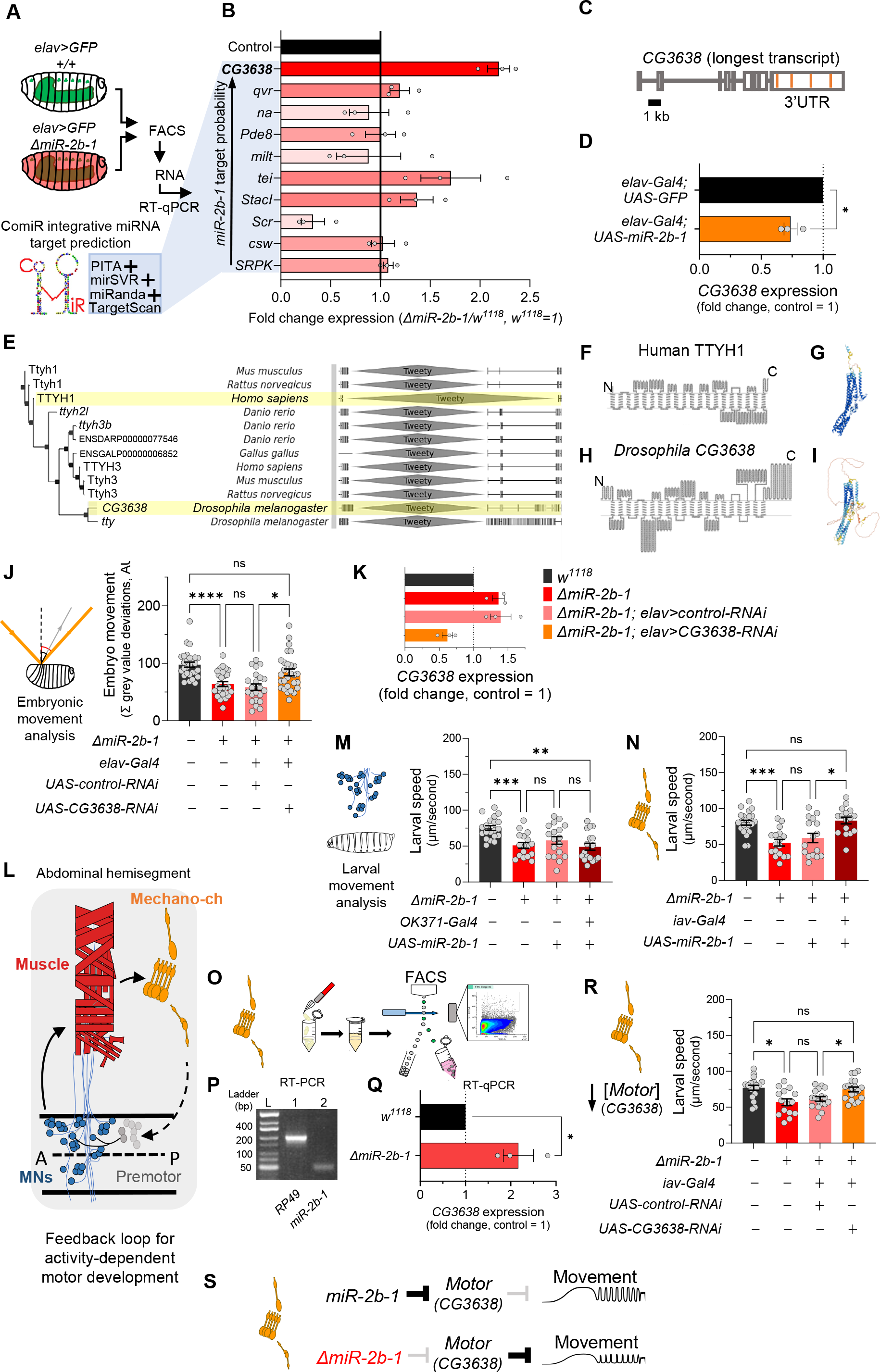
The genetic and cellular mechanisms that link miR-2b-1 to embryonic movement (**A**) Workflow for the FACS and RT-qPCR experiments shown in panel B and schematic describing the ComiR miRNA target prediction tool used to generate the list of candidate *miR-2b-1* targets. (**B**) Expression analysis (qPCR) of 10 predicted *miR-2b-1* target genes shown as fold change between *ΔmiR-2b-1* mutant and control *w^1118^* embryos (three biological replicates). Targets are listed from top to bottom by descending probability score. [The black control bar, set to 1, represents expression of each gene in control *w^1118^* embryos]. Note that upregulation of *CG3638* is statistically significant (p=0.0169). (**C**) Schematic of the *CG3638* transcript with *miR-2b-1* target sites indicated (orange lines). (**D**) Whole embryo qPCR experiment showing a reduction of *CG3638* expression in *elav- Gal4>UAS-miR-2b-1* embryos (orange bar), relative to control *elav-Gal4>UAS-GFP* embryos (black bar). (**E**) Evolutionary conservation of the *CG3638* protein across a wide range of invertebrate and vertebrate species (left), as determined with PhylomeDB 5 software (Huerta-Cepas *et al*., 2014) [*Homo sapiens* and *Drosophila melanogaster* highlighted in yellow]. Gene schematics highlighting the conserved Tweety domain are shown on the right. (**F-I**) Transmembrane domain structure (left) and AlphaFold structural predictions (right) for Human TTYH1 (F-G) and *Drosophila CG3638* (H-I). (**J**) Embryonic movement quantification (summation of MGV deviations) of *ΔmiR-2b-1*, *elav>CG3638-RNAi* embryos (orange bar) during the neurogenic phase compared to control *w^1118^* (black bar), *ΔmiR-2b-1* mutant (bright red bar) and control *ΔmiR-2b-1*, *elav>control-RNAi* embryos (faded red bar). (**K**) qPCR expression profiling of *CG3638* in whole embryos of the genotypes tested in panel J. (**L**) Diagram describing key cell types that form a feedback loop for activity-dependent motor development. Motor neurons (MNs, blue) induce muscle (red) movements which are in turn detected by proprioceptive chordotonal organs (Mechano-ch, orange) and feed-back into the CNS to regulate activity patterns. (**M-N**) Reconstitution experiments that restore *miR-2b-1* expression in specific cellular elements related to embryonic movement circuitry. (M) Quantification of larval speed in control *w^1118^*(black); *ΔmiR-2b-1* mutant (red); *ΔmiR-2b-1*, *UAS-miR-2b-1* parental control (pink) and *ΔmiR-2b-1*, *OK371-Gal4>UAS-miR-2b-1* experimental embryos (brown). (N) Quantification of larval speed in *ΔmiR-2b-1*, *iav-Gal4>UAS-miR-2b-1* (brown) and control genotypes as in panel M. (**O**) Schematic describing FACS isolation of embryonic chordotonal organs. (**P**) Mature *miR-2b-1* (right) is expressed in chordotonal organs isolated during the neurogenic phase (RP49 expression shown on left). (**Q**) Chordotonal specific qPCR expression profiling of *CG3638* in *ΔmiR-2b-1* mutant and control *w^1118^* embryos. (**R**) Average larval speed of *ΔmiR-2b-1*, *elav>CG3638-RNAi* (orange) compared to control *w^1118^* (black), *ΔmiR-2b-1* mutant (red) and control *ΔmiR-2b-1*, *UAS-control-RNAi* (pink). (**S**) Model for the mechanism by which *miR-2b-1* acts to control embryonic movement in chordotonal organs. Under normal (control) conditions (top), *miR- 2b-1* inhibits the expression of *CG3638* and thereby enables normal movement. In *ΔmiR-2b-1* mutants (bottom), de-repression of *CG3638* expression leads to a reduction in embryonic movement.

A genuine genetic target for a given miRNA is predicted to: (i) be de-repressed (up-regulated) in the absence of the miRNA; and (ii) be down-regulated under miRNA ectopic expression. Analysis of *CG3638* expression shows that this target meets the predictions of a genuine *miR-2b-1* target in full: expression of *CG3638* is upregulated in *ΔmiR-2b-1* mutants (Figure 4B) and reduced under neural over-expression of *miR-2b-1* (Figure 4D). As mentioned above, phylogenetic analysis of *CG3638* reveals that it belongs to an evolutionarily conserved family of chloride channel genes[41, 42], with representatives across distantly related lineages of insects and vertebrates including mammals, strongly indicating a functional role (Figure 4E). Comparison of the properties of *CG3638* and its human orthologue reveal the characteristic seven trans-membrane domains with an external carboxyterminal topology (Figure 4F, 4H) further supporting orthology, and applying *AlphaFold* – an artificial intelligence computational method able to predict protein structures with atomic accuracy [43] – to the proteins encoded by the *Drosophila CG3638* gene and the human TTYH1 gene reveals the marked similarities between these two polypeptides (Figure 4G, 4I).

To explore the relationship between *CG3638* and the embryonic movement phenotype displayed by *ΔmiR-2b-1* mutants, we tested the effects of an artificial reduction of *CG3638* in the genetic background of the miRNA mutant (Figure 4K). In this scenario, should the levels of expression of *CG3638* be relevant to the triggering of the embryonic movement phenotype, we predict that a reduction in *CG3638* expression levels should compensate its cellular effects, and, accordingly, reduce or even rescue the embryonic phenotype. The results of this experiment show that this is indeed the case, with embryonic movement of the *ΔmiR-2b-1* mutant effectively rescued by a reduction in *CG3638* (Figure 4J). Based on its modulatory role in embryonic movement we termed *CG3638* as *<u>Mo</u>vement modula<u>tor</u> (Motor)*.

### Mapping the focus of action of miR-2b-1 within the known networks underlying embryonic movement

Having observed the effects of *miR-2b-1* on embryonic movement we wondered about the site of action of this miRNA in regard to circuit components previously linked to embryonic movement. In this respect, previous work has identified embryonic motor neuronal components, as well as interneurons and elements of the sensory system as playing key roles in the control of motor development. These include a motor component that includes all embryonic motor neurons which command the stereotypic array of muscles in the embryonic body wall [44, 45], as well as specific elements of the sensory system – in particular the chordotonal system – which detect early myogenic movements in the embryo and transmit the information to the pattern generators thus modulating motor patterns (Figure 4L) [17, 19, 46, 47]. To determine which one of these known circuitry elements might be the principal focus of action of *miR-2b-1* in connection to embryonic movement control we artificially expressed *miR-2b-1* in each motor and sensory circuit elements – using drivers *OK371-Gal4* [48] and *iav-Gal4* [49], respectively – in an otherwise null mutant background for *miR- 2b-1*, asking whether these genetic restorations were sufficient to improve or perhaps even rescue normal movement patterns. To ensure that circuit-specific Gal-4 drivers were active and UAS-driven miRNA levels achieved the necessary cumulative values for biological activity we chose to measure effects on early larval movement patterns tested in 30-min old first instar larvae (L1s). The results of these experiments are shown in Figure 4M-N, S3A-B. Here we observe that restoring expression of *miR-2b-1* in the motor neuronal domain defined by the OK371 driver is insufficient to affect the defective movement patterns observed in *miR-2b-1* null mutants (Figure 1B-C and Figure 4M, S3A). In contrast, re-establishing expression of the miRNA in all eight chordotonal organs leads to a full rescue of the motor phenotype (Figure 4N, S3B) suggesting that this aspect of the sensory system might be the one where *miR-2b-1* exerts relevant actions during the normal development of movement. In line with this, we observe that the mature *miR-2b-1* transcript is indeed expressed in FACS-isolated embryonic chordotonal organs prepared from wild type embryos (Figure 4P) and, that the genetic target of *miR-2b-1*, *Motor*, is also expressed in these cells in normal embryos (Figure 4Q, top). In addition, expression of *Motor* in chordotonal organs prepared from *miR-2b-1* null mutants is up-regulated (Figure 4Q, bottom) lending further support to a model in which *Motor* is de-repressed in these specific sensory elements. Furthermore, artificial reduction of *Motor* implemented as a stratagem to decrease the effects of de-repression specifically within the chordotonal system is sufficient to rescue normal movement patterns (Figure 4R). Altogether, these findings strongly suggest that *miR-2b-1* impacts the emergence of embryonic movement, at least in part, via effects on the sensory circuit components that underlie motor development, rather than affecting the actual generation of motor patterns.

Our work identifies a miRNA system that plays a role in the emergence of embryonic movement in the fly embryo, and offers a new approach to analyse the roles of non-coding RNAs and protein coding genes at the critical period when patterned movement develops. Understanding the molecular elements controlling the onset of motor development in *Drosophila* will put us one step closer to understanding the molecular basis of embryonic movement in other species, including vertebrates, whose embryos seem to undergo remarkably similar transition phases to those reported here [20].

## ACKNOWLEDGEMENTS

We wish to thank all members of the Alonso Lab for helpful discussions and feedback on this work. We are also grateful to Marta Zlatic for sharing Drosophila lines, and to the Bloomington Stock Center and the Vienna *Drosophila* Resource Center for providing fly stocks. This research was funded by a Wellcome Trust Investigator Award (098410/Z/12/Z) made to C.R.A., a Wellcome Trust Investigator Award (220277/Z20/Z) made to T.B. and a UK Medical Research Council Project Grant (Ref: MR/S011609/1) given to C.R.A.

