## Supplementary Information for "A microRNA that controls the emergence of embryonic movement"


This file contains the following information:

- Supplementary Figures 1-4
- Movie Legend for Movie S1
- Materials and Methods
- References

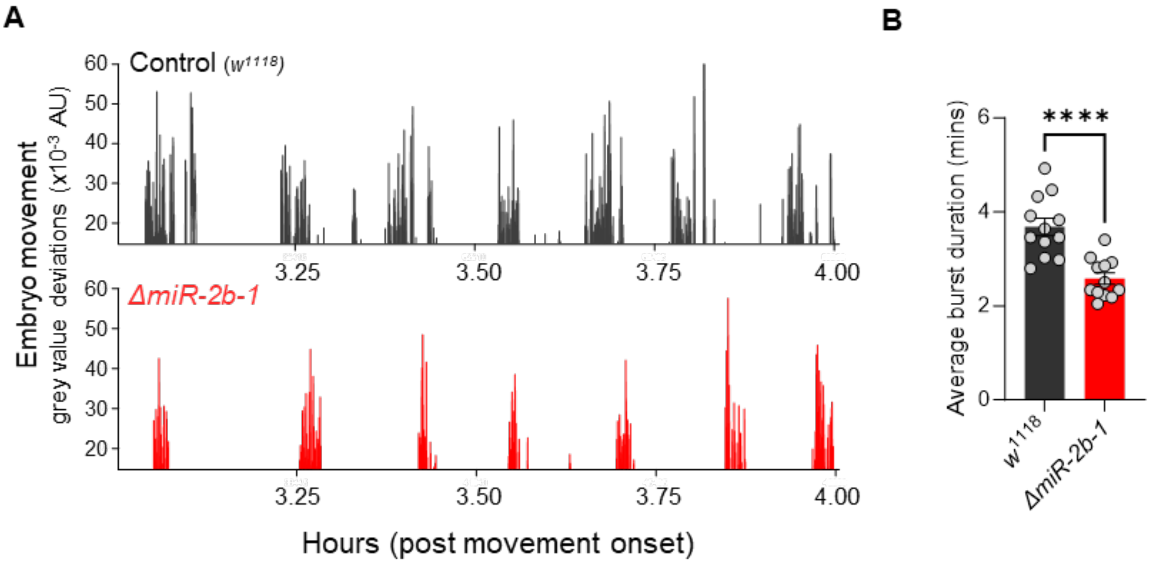


**Figure S1.** ***miR-2b-1*** **decreases movement burst duration during the neurogenic phase**
(**A**) Representative movement bursts from control (*w*$^{1118}$, top) and *ΔmiR-2b-1* (bottom) embryos.
(**B**) Quantification of average burst length (minutes) during the neurogenic phase of movement in control (*w*$^{1118}$, black) and *ΔmiR-2b-1* (red) animals.

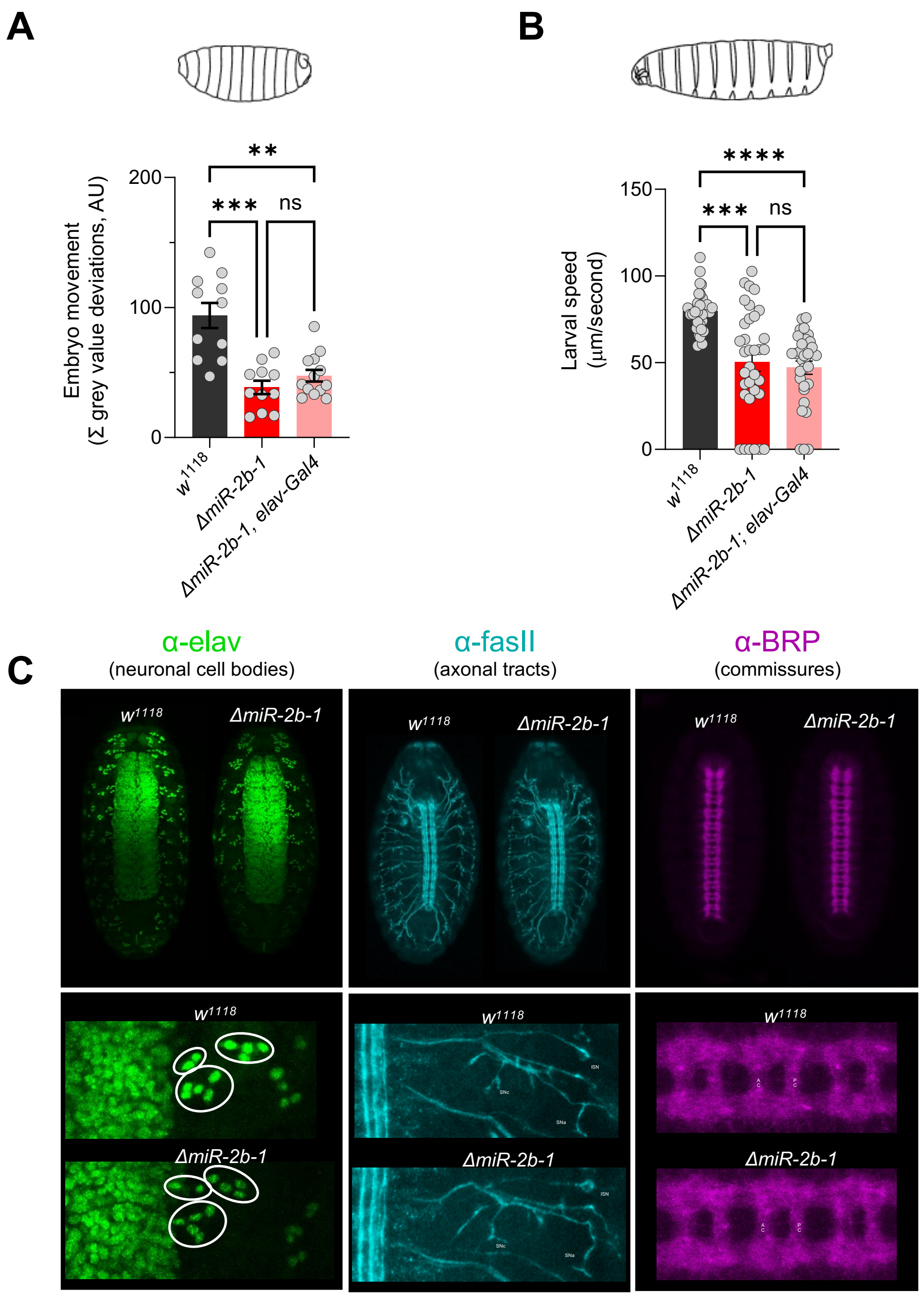


**Figure S2. Additional controls for *miR-2b-1* genetic reconstitution in neurons; structural analysis in *ΔmiR-2b-1* embryos**
(**A**) Summation of MGV deviations during the neurogenic phase of embryonic movement in control $w^{1118}$ embryos (black bar); *ΔmiR-2b-1* mutant embryos (bright red bar); *ΔmiR-2b-*, *elav-Gal4* pan-neuronal parental control embryos (faded red bar).
(**B**) Average L1 larval speed for the same genotypes used in the embryonic genetic reconstitution experiment shown in panel (A).
(**C**) Immunohistochemical analyses of $w^{1118}$ and *ΔmiR-2b-1* embryonic nervous system structures. Zoomed in images for one hemisegment are shown in the lower panels.

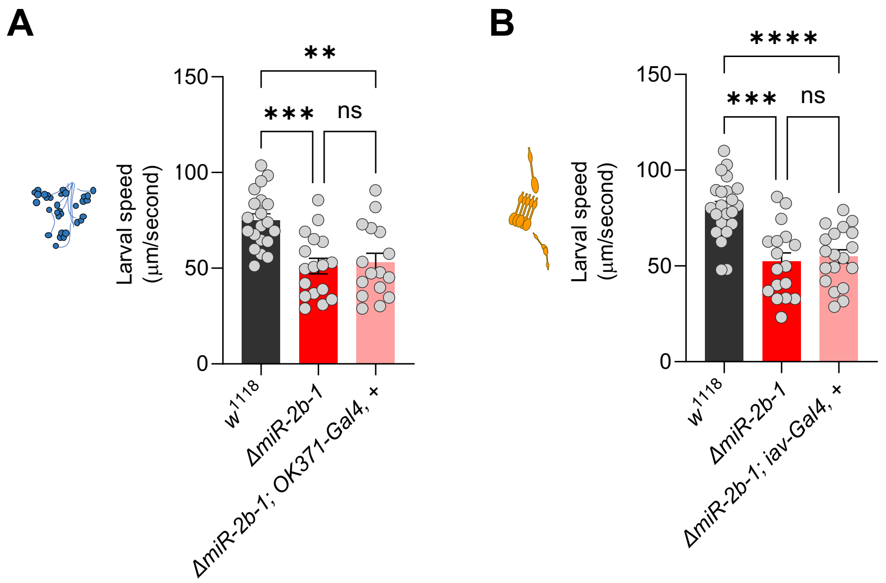


**Figure S3. Additional controls for genetic reconstitutions in motor neurons and chordotonal organs**

(**A**) Average larval speed of control $w^{1118}$ larvae (black bar); *ΔmiR-2b-1* mutant larvae (bright red bar); *ΔmiR-2b-*, *OK371-Gal4* motor neuron parental control larvae (faded red bar).

(**B**) Average larval speed of control $w^{1118}$ larvae (black bar); *ΔmiR-2b-1* mutant larvae (bright red bar); *ΔmiR-2b-*, *iav-Gal4* chordotonal organ parental control larvae (faded red bar).

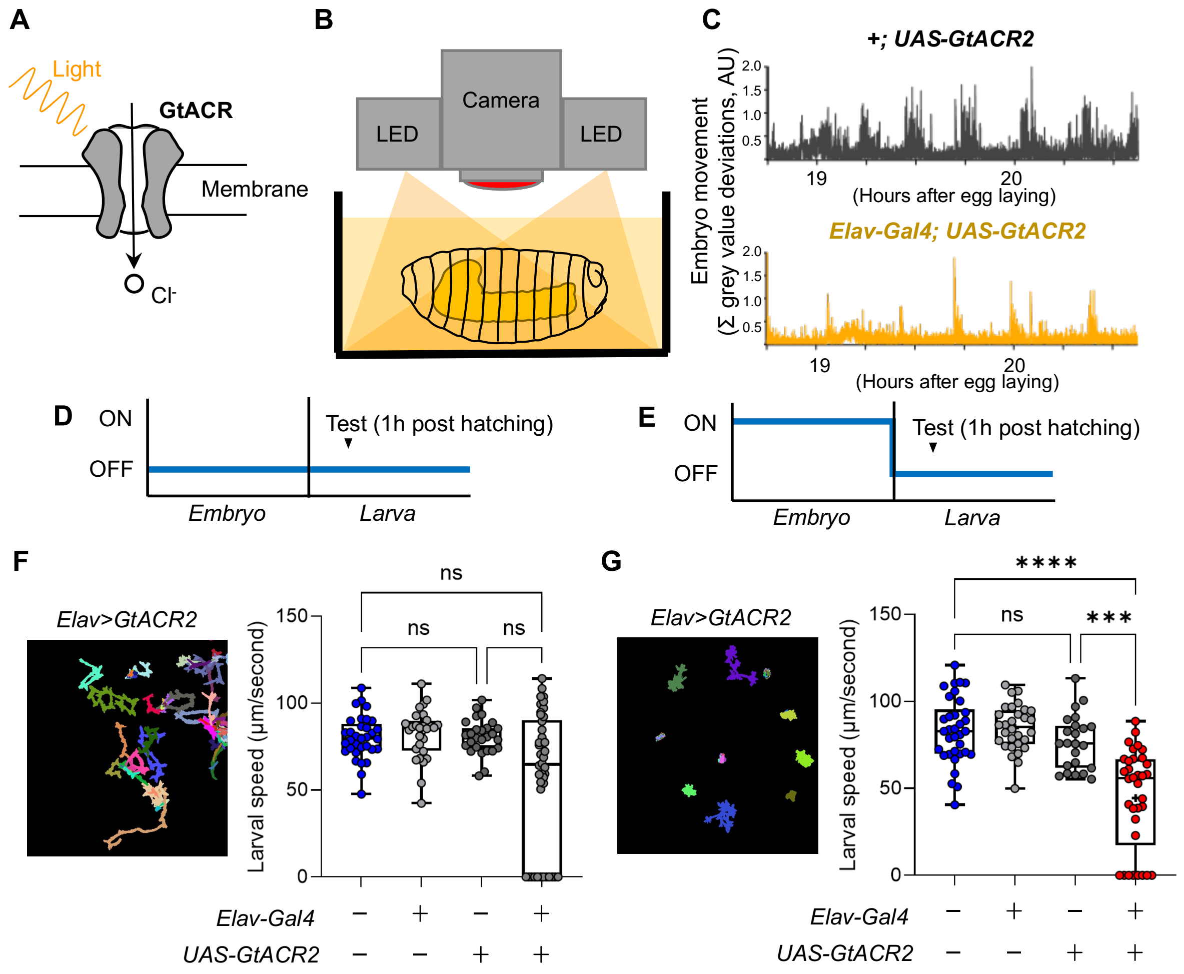


**Figure S4. Inhibiting neural activity during embryogenesis reduces larval locomotor speed**

(**A**) Graphic showing GtACR2 channel gating chloride ions across the cell membrane in response to photon stimulation. (**B**) Diagram describing our experimental setup, with embryos illuminated by red light throughout embryogenesis and (**C**) Embryo 'movement over time' traces for the control *(+;UAS-GtACR2*) and experimental (*elav-Gal4; UAS-GtACR2*) genotypes in response to red light stimulation. (**D-E**) Schematics showing the time course of red light stimulation. NB: ON leading to activation of GtACR2 and repression of neural activity; OFF leading to inactivation of GtACR2 (no effect on neural activity). (D) control experiment; (E) inhibition experiment. Developmental time is shown on the x axis. (**F-G**) Larval movement analysis using the FIM platform. (F) Control experiment (see D). Left: FIM cumulative larval tracks of *elav-Gal4>UAS-GtACR2* L1 larvae in the control condition (no inhibition, see D); Right: quantification of larval locomotor speed across a range of genotypes (see bottom of graph). (G) Experimental condition (see E). Left and right panels show larval tracks and quantification data respectively, as in panel F.

### MOVIE LEGENDS

**Movie S1. Embryonic movement**
Movement from a control $w^{1118}$ embryo recorded across the movement period from 16 to 21 hours after egg laying, 300X speed.

### MATERIALS AND METHODS

| REAGENT or RESOURCE | SOURCE | IDENTIFIER |
|---|---|---|
| *Drosophila* strains | | |
| *w[1118]* | BDSC | 5905 |
| *w[1118]; TI{w[+mW.hs]=TI} mir-2b-1[KO]* | BDSC | 58915 |
| *w[1118]; P{w[+mC]=GAL4-elav.L}3* | BDSC | 458 |
| *w[1118]; P{w[+mW.hs]=GawB}VGlut[OK371]* | BDSC | 26160 |
| *w[*]; P{w[+mC]=iav-GAL4.K}3* | BDSC | 52273 |
| *UAS-Kir* | Bate Lab, Cambridge (Baines et al., 2001) | N/A |
| *w[1118]; P{y[+t7.7] w[+mC]=10XUAS-IVS-myr::GFP}attP2* | BDSC | 32197 |
| *w[1118]; P{y[+t7.7] w[+mC]=UAS-LUC-mir-2b-1.T}attP2* | BDSC | 41128 |
| *w[1118]; UAS-IVS-Syn21-GCaMP6s-P2A-nls-tdTomato-p10 (JK66B)* | Zlatic Lab, Cambridge | N/A |
| *UAS 40D RNAi-KK* | VDRC | 60101 |
| *CG3638 RNAi-KK* | VDRC | 102444 |
| *UAS-GtACR2* | BDSC | 92984 |
| Primers | | |
| In-situ hybridisation control sense probe F: ATTTAGGTGACACTATAGAGAATTCAACGCGCAGCATC | Sigma-Aldrich | N/A |
| In-situ hybridisation control sense probe R: ACACCAAACTGTCCCGATCC | Sigma-Aldrich | N/A |
| In-situ hybridisation experimental Btk anti-sense probe F: AGAATTCAACGCGCAGCATC | Sigma-Aldrich | N/A |
| In-situ hybridisation experimental Btk anti-sense probe R: ATTTAGGTGACACTATAGACACCAAACTGTCCCGATCC | Sigma-Aldrich | N/A |

| | | |
|---|---|---|
| miRNA PCR RT mix primer 1 | Sigma-Aldrich | N/A |
| miRNA PCR RT mix primer 2 | Sigma-Aldrich | N/A |
| miRNA PCR RT mix primer 3 | Sigma-Aldrich | N/A |
| miRNA PCR RT mix primer 4 | Sigma-Aldrich | N/A |
| miRNA PCR RT mix primer 5 | Sigma-Aldrich | N/A |
| miRNA PCR RT mix primer 6 | Sigma-Aldrich | N/A |
| miRNA PCR RT mix primer 7 | Sigma-Aldrich | N/A |
| miRNA PCR RT mix primer 8 | Sigma-Aldrich | N/A |
| miRNA PCR RT mix primer 9 | Sigma-Aldrich | N/A |
| miRNA PCR RT mix primer 10 | Sigma-Aldrich | N/A |
| miRNA PCR RT mix primer 11 | Sigma-Aldrich | N/A |
| miRNA PCR RT mix primer 12 | Sigma-Aldrich | N/A |
| PCR RP49 (RpL32) F: CCAGTCGGATCGATATGCTAA | Sigma-Aldrich | N/A |
| PCR RP49 (RpL32) R: TCTGCATGAGCAGGACCTC | Sigma-Aldrich | N/A |
| PCR/qPCR miR-2b-1-5p F: GGTCTTCAAAGTGGCAGTG | Sigma-Aldrich | N/A |
| PCR/qPCR miR-2b-1-5p R: GTCCAGTTTTTTTTTTTTTTTCATGTC | Sigma-Aldrich | N/A |
| PCR/qPCR miR-2b-3p F: AGTATCACAGCCAGCTTTG | Sigma-Aldrich | N/A |
| PCR/qPCR miR-2b-3p R: GTCCAGTTTTTTTTTTTTTTTGCTC | Sigma-Aldrich | N/A |
| qPCR CG3638_PP20655 F: TCCTTGGTCATCATTACGCTGA | Sigma-Aldrich | N/A |
| qPCR CG3638_PP20655 R: CCATTATGGAAATCATCGTTGCC | Sigma-Aldrich | N/A |
| qPCR qvr_PP25844 F: CCTTTCAACTATACAGCCCTGC | Sigma-Aldrich | N/A |
| qPCR qvr_PP25844 R: TGTAACTGTGACGTACACATGC | Sigma-Aldrich | N/A |
| qPCR na_PP34188 F: ACCTTTCCTCGCGGATTACG | Sigma-Aldrich | N/A |
| qPCR na_PP34188 R: CCACAGCTTGTTCACCCAC | Sigma-Aldrich | N/A |
| qPCR Pde8_PP11165 F: CCGAGAAAATCCGTCCAGC | Sigma-Aldrich | N/A |
| qPCR Pde8_PP11165 R: CAGCGGTCTTGGTCTTTCATTA | Sigma-Aldrich | N/A |
| qPCR milt_PP21284 F: GCAGACGATGGCACAGATACT | Sigma-Aldrich | N/A |
| qPCR milt_PP21284 R: CGTCGAGCAGGGAGTTGAC | Sigma-Aldrich | N/A |
| qPCR CG17716_PP26416 F: GTCCGTGGTCTATGCGGAG | Sigma-Aldrich | N/A |
| qPCR CG17716_PP26416 R: ATGAAGCGATAGTCGGTGACG | Sigma-Aldrich | N/A |

| | | |
|---|---|---|
| qPCR Stacl_PP10900 F: GCTGCGTCCCAATCTGGAT | Sigma-Aldrich | N/A |
| qPCR Stacl_PP10900 R: CGTGTGTGCCCTCTCAGAAT | Sigma-Aldrich | N/A |
| qPCR Scr_PP19886 F: GGCGGCCTATACGCCTAAC | Sigma-Aldrich | N/A |
| qPCR Scr_PP19886 R: CGGCTGTAGCTGCGTGTAG | Sigma-Aldrich | N/A |
| qPCR csw_PP8739 F: TTTGGCACCTTGTCGGAACT | Sigma-Aldrich | N/A |
| qPCR csw_PP8739 R: CCAGAAACCTCCCTTGACCAG | Sigma-Aldrich | N/A |
| qPCR SRPK_PA60244 F: ATCCGCTGACTGAGGGCACTG | Sigma-Aldrich | N/A |
| qPCR SRPK_PA60244 R: GTAGAGTTTTCCAGTTGTGG | Sigma-Aldrich | N/A |

**Experimental model details**

*Drosophila melanogaster* fruit flies were maintained by standard means; in 25°C incubators with 50-60% humidity; on a 12-hour light/dark cycle; with molasses food. See reagent and resource table for all *Drosophila* strains used in this project and the respective sources.

**Collection of samples for behavioural experiments**

Flies were kept at 25˚C in collection cages with food plates consisting of apple juice agar and yeast paste. Embryos were collected by placing a fresh food plate in the collection cage and allowing flies to lay eggs for 1 hour. Prior to all embryo collections, a pre-collection of 1 hour was performed to reduce female egg storage. In experiments where some embryos were of genotypes that included GFP-tagged balancer chromosomes, those individuals were selected against by fluorescence microscopy. Selected embryos were gently moved to a fresh plate and allowed to develop at 25˚C. All genotypes underwent selection by fluorescence microscopy to ensure consistent exposure to light across groups compared.

### Embryo chamber design and 3D-printing

The 3D-printed embryo chamber was designed on paper to dimensions of 45mm (L) X 15mm (W) X 3mm (D). Four sub-chambers were designed within the main chamber, each with dimensions of 5mm (L) X 5mm (W) X 0.5mm (D) and divided by a boundary wall of 0.4mm. The design was subsequently coded in OpenSCAD software and printed on a Formlabs Form 2 desktop 3D-printer using biocompatible BioMed Black resin.

### Embryo movement experiments

Embryo collections were aged to 14 (±0.5) hours after egg laying (AEL) prior to selection of individuals with the correct genotype determined via the fluorescence balancer. Embryos were subsequently adhered to a piece of tesa® double-sided tape that itself was adhered to a microscope slide. Using one end of a pair of dissection forceps and observing through a brightfield microscope, embryos were gently rolled on the tape to break them out of the egg chorion before being transferred to a well of an embryo chamber previously glued with tesa® glue dissolved in heptane. 6 embryos were transferred one-by-one to the well prior to the addition of 3 μl of Halocarbon oil (a 50:50 mix of series 27 and series 700). All manipulations, from the dechorionation of the first embryo to the addition of Halocarbon oil, were done within 3 minutes to ensure minimal dehydration of embryos. This process was repeated for each of the remaining chamber wells.

### Embryo movement recording

Movements of embryos across all 4 wells of the embryo chamber were recorded simultaneously using a Leica DFC 340 FX camera mounted on a Leica M165 FC microscope, with a resolution of 480 x 360 pixels and a frame rate of 4 frames per second. Incident lighting was directed laterally onto the embryo chamber to avoid glare to the camera from the surface of the Halocarbon oil. Consistent lighting conditions across the 4 wells of the embryo chamber were ensured through measurement of pixel intensities (mean grey values) within each recorded well in ImageJ software. Recordings were carried out for at least 10 hours to capture the entire duration of embryonic movement up until larval hatching and files were stored in the AVI format with MJPEG compression to ensure compatibility with downstream analysis software. All recordings were carried out at 25˚C.

**Embryo movement analysis**

AVI files were opened in ImageJ software and a rectangular ROI of consistent size was applied over each embryo within the chamber. The ‘RoiSet’ of up to 24 ROIs was saved and then used to ‘multi-measure’ the mean grey value (MGV) of each ROI for each frame of the recording – rapid changes within which were caused by embryo movements that altered reflected light to the camera. The resulting list of MGV was exported to Excel software where a background subtraction was applied to remove slow-scale changes to MGV that occurred due to gradual changes in embryo morphology. This involved the generation of a moving average for the MGV of each embryo with a sliding-window of 60 frames or 15 seconds, which was then subtracted from the MGV for each frame. The choice of 15 seconds was made empirically based on the duration of individual movements and the rate of morphological change, particularly tracheal gas-filling. Absolute values were taken for deviations from the baseline to create traces that represent embryo movements over time and for all quantifications. Traces were subsequently cropped at larval hatching based on when the vitelline membrane was breached by the head of the larva. Movement was quantified by summing deviations in MGV from baseline prior to larval hatching. A threshold value of 0.01 MGV deviations from baseline, determined empirically by the comparison of traces from unfertilised and live embryos and found to be applicable across recordings, was applied to filter noise that was unrelated to embryo movement. Traces where a different larva had entered the ROI following hatching were removed from the analysis. Fast Fourier Transform (FFT) analysis was performed in Igor PRO software and was applied to 1-hour overlapping (30-minute overlaps) sliding windows of movement traces to extract information about the frequency spectrum of movements.

**Larval movement experiments**

For all larval movement experiments, we used an imaging method based on frustrated total internal reflection (FTIR) – known as FIM (FTIR-based Imaging Method). This allowed for the quantification of larval movement with a high degree of consistency and accuracy. See Risse *et al*. (2013; 2017) for more information. A FIM table was obtained from the University of Münster, department of Computer Vision and Machine Learning Systems, for this purpose. First instar larvae were gently moved to fresh agar plates for assessment on the FIM table within 30 minutes of hatching to ensure consistency of age across larvae tested and reduce the possibility of differences in motor learning. At least 25 larvae were assessed for each genotype across 3 independent recordings. TIFF images were captured for 3 minutes at a resolution of 1200x1200 pixels and frame rate of 7 frames per second using a Basler acA2040-90um camera. All recordings were carried out at 25˚C in low-light conditions.

**Larval movement analysis**

TIFFs were opened in FIMTrack software (Risse *et al.*, 2017) before running the tracking algorithm with the 'minimum larval size' set to 40. All other settings were left as default. Partial tracks, due to larvae crawling off the plate or into one another, were removed from the analysis. Larvae that did not move were considered a 0 value as the FIMTrack software was unable identify them. The L1_acc_dis parameter was extracted for each larva and this was taken as a quantitative measure of larval movement and compared across genotypes.

**Statistical analyses**

All statistical analyses were performed in GraphPad Prism software. The normality of each dataset was determined by the agreement of four tests: D'Agostino & Pearson; Anderson-Darling; Shapiro-Wilk; Kolmogorov-Smirnov. Datasets that at least one of these normality tests identified as not having a normal distribution were further assessed by nonparametric tests. Multiple Mann-Whitney tests with Bonferroni correction were used for comparison of two genotypes in the myogenic and neurogenic phases or at the larval stage. The parametric Brown-Forsythe and Welch ANOVA with Dunnett's T3 multiple comparisons tests, or nonparametric Kruskal-Wallis ANOVA with Dunn's multiple comparisons tests were used for comparisons of more than two genotypes assessed in parallel during the miRNA rescue and RNAi experiments. **** = $p<0.0001$, *** = $p<0.001$, ** = $p<0.01$, * = $p<0.05$.

**In-situ hybridisation**

In-situ hybridisation probes were designed to be 500-1000 bases in length and complementary to the exons of Btk mRNA. A negative control probe made in the sense orientation to the target mRNA was used alongside the experimental anti-sense probe, at the same concentration, to control for non-specific binding. See reagent and resource table for all primers used in probe synthesis. An SP6 polymerase tag was added to the forward (sense probe) or reverse (anti-sense probe) primer for transcription. *$w^{1118}$* embryos were pre-hybridised in hybridisation solution (50% formamide) for at least 2 hours prior to overnight hybridisation at 55˚C. Post-hybridisation, embryos were blocked and stained with an α-DIG-POD antibody (Roche 11207733910), prior to a fluorescein tyramide treatment to increase signal strength and imaging with a confocal fluorescence microscope.

**Fluorescence activated cell sorting (FACS)**

For cell dissociation, embryos were collected and aged to 18 (±0.5) hours AEL prior to dechorionation and digestion in a haemolymph-like solution (90 mM NaCl, 25 mM KCl, 10 mM HEPES, 80 mM D-glucose, 4.8 mM NaHCO3, pH 7) with 0.25% trypsin at 37˚C and gentle mechanical disruption. The cell solution was passed through a 40 nm filter immediately prior to sorting. Cell sorting was performed on a BD FACSmelody cell sorter (BD Biosciences) calibrated to sort GFP+ cells by sorting 100,000 cells from embryos of the *UAS-GFP* genotype (without a Gal4 driver) and observing the highest level of fluorescence seen from these cells, before gating the cell sorter to only isolate cells with a level of fluorescence above this. For each biological replicate of each genotype, 10,000 cells were sorted into 470 µl TRIzol reagent (Invitrogen) for downstream RNA extraction.

**Conventional and real-time quantitative PCR**

Conventional PCR was performed with standard Taq DNA polymerase (New England Biolabs – M0273)  For all reactions, 30 amplification cycles were run with 0.4 µM final concentration of each primer (see reagent and resource table for a list of all primers used) and a 60˚C annealing temperature. qPCR reactions were performed with LightCycler SYBR Green I reagents (Roche – 04707516001). For all reactions, 40 amplification cycles were run with 0.25 µM final concentration of each primer (see reagent and resource table for all primers used) and a 60˚C annealing temperature. All reactions were run with 2 technical replicates and any groups compared in downstream analysis were run on the same reaction plate. Continuous melt curves were examined to assess whether a single amplicon was amplified by each primer set and no-template controls were also run to confirm a lack of primer-dimer formation. Primer efficiency was determined by a standard curve of 6 cDNA dilutions and only those with efficiencies between 1.9 and 2.2 were used. Efficiency – $E$ – was calculated with the following equation (Pfaffl, 2001):

$$E = d^{-1/-s}$$

Where *d* is the dilution factor and *s* is the slope of the curve. Fold change in transcript expression between two experimental conditions was calculated using $C_T$ values obtained from the qPCR experiment with the following equation (Pfaffl, 2001):

$$Fold\ change = \frac{2^{CT\ gene\ of\ interest\ (control-mutant)}}{2^{CT\ reference\ gene\ (control-mutant)}}$$

**Mature miRNA PCR**

PCR to specifically amplify mature miRNA transcripts utilised a protocol based on a single reverse transcription reaction for all microRNAs combined with PCR using two, mature miRNA-specific DNA primers (Balcells *et al.*, 2011). Poly(A) tailing of total RNA prior to reverse transcription ensured that miRNA transcripts would be included in the cDNA product, due to addition of a poly(A) tail to each. Reverse transcription was performed with a modified oligo (dT) primer that included a 5' universal tag (see reagent and resource table for primers used). This 5' universal tag enabled mature miRNA-specific primer sets to bind in downstream PCR experiments. Primer sets for different miRNAs were designed in miRprimer software (Busk, 2014) with specificity and efficiency was confirmed by dilution series and melt curve analyses, in addition to running the primer sets with cDNA from miRNA mutant samples to confirm a lack of amplification. Both conventional PCR and qPCR were used for miRNA specific PCR experiments.

**Calcium imaging**

Calcium imaging was performed using a Leica DM6000 epifluorescence microscope with a 10X objective. Embryos were aged to 14hAEL, dechorionated and adhered in the ventral-up orientation to a clear glass microscopy slide using Tesa tape glue. Images were captured sequentially using an ET470 40x ET525 50m band pass filter for detecting GCaMP6s signal and a ET545 25x ET605 70m set for tdTomato, with a single image cycle occurring over 1.5 seconds. Recording was performed for 6 hours or until hatching occurred. Fluorescence signals from each channel were measured in FIJI software using ROIs of equal size placed over each embryo. To calculate $\Delta F/F$, the GCaMP6s reading for each frame was divided by the tdTomato reading for the same frame, before subtraction of the baseline calculated as the minimum value of this ratio in a 10-minute window centred around each frame.

**Bioinformatic miRNA target prediction**

Potential miRNA targets were predicted with the bioinformatic combinatorial target prediction tool ComiR (Coronnello *et al.*, 2013) that draws upon weighted prediction scores from four major miRNA target prediction algorithms- miRanda, PITA, TargetScan and mirSVR - before integrating them through a machine learning model trained on biochemical data for miRNA-mRNA interactions (*Drosophila* AGO1 IP data – Hong *et al.*, 2009). From this, a list of predicted targets for a miRNA was generated and ranked by probability score. The following two criteria were applied to filter for targets with a probable role in nervous system functional development: (i) at least one of the following major GO terms: receptor; receptor binding; transporter; small molecule binding; development; nervous system process; behaviour. (ii) Embryonic expression according to modENCODE Development RNA-Seq. A filtered list of the top-10 predicted targets of miR-2b-1, sorted by ComiR score, was subsequently obtained for assessment in biochemical experiments.

**Bioinformatic analysis of CG3638**

Evolutionary conservation of *CG3638* protein was determined with PhylomeDB 5 software (Huerta-Cepas *et al*., 2014). For structural analysis, AlphaFold (Jumper *et al*., 2021) software was used to predict protein structure and SACS MEMSAT2 (Jones *et al*., 1994) software was used to visualise transmembrane domains.

**Optogenetic inhibition of neural activity**

Embryos were placed on plain 1.5% agar plates and exposed to red light – at 650nm wavelength and 5,000 lux as measured on a EXTECH Instruments 401020 lux meter – for 21 hours. Embryos were subsequently moved to a dark room until hatching, checked regularly under brief weak red light (<1000 lux). Once larvae had hatched, they were moved to a different plan 1.5% agar plate and left for 1 hour under dark conditions. Subsequently, the agar plate was placed on a FIM table and locomotion was tracked for 3 minutes.
